# Aberrant expression of the COX2/PGE_2_ axis is induced by activation of the RAF/MEK/ERK pathway in BRAF^V595E^ canine urothelial carcinoma

**DOI:** 10.1101/786095

**Authors:** R. Yoshitake, K. Saeki, S. Eto, M. Shinada, R. Nakano, H. Sugiya, Y. Endo, N. Fujita, R. Nishimura, T. Nakagawa

## Abstract

Cancer-promoting inflammation is an important event in cancer development. The cyclooxygenase 2 (COX2)/prostaglandin E2 (PGE_2_) axis is a prominent inducer of inflammation. Canine urothelial carcinoma (cUC) uniquely overexpresses PGE_2_ and is often managed well with COX inhibitor monotherapy. In most cases, cUC tissue harbours homologous human BRAF^V600E^ mutation, which causes aberrant activation of the RAF/MEK/ERK pathway in human cancer. However, mechanisms underlying aberrant PGE_2_ production and the importance of the *BRAF* mutation remain unclear. We show that activation of the RAF/MEK/ERK pathway in *BRAF* mutant cUC cells leads to COX2 overexpression and PGE_2_ production. Drug screening revealed that treatment with inhibitors of the arachidonic acid cascade (FDR<0.086), RAF/MEK/ERK pathway (FDR<0.067), and p38/JNK pathway (FDR<0.067) significantly reduced PGE_2_ production in cUC cells. We also validated the association between RAF/MEK/ERK pathway activation and COX2/PGE_2_ overexpression in *BRAF* mutant cUC cells using protein detection techniques. In histochemical analysis, *BRAF* mutant cUC tissue showed higher COX2 expression. Therefore, the driver mutation in the *BRAF* gene probably promotes tumour-promoting inflammation. These findings would benefit dogs suffering from cUC and can be extrapolated to human cancer. Finally, cUC can serve as a valuable model to elucidate the association between driver mutations and tumour-promoting inflammation.

## Introduction

Inflammation occurring in cancer tissues promotes cancer progression by providing various factors, such as growth factors, pro-angiogenic factors, enzymes required for cancer invasion and metastasis, and immune-suppressive factors. Thus, such inflammation is called cancer-promoting inflammation^1, 2^. Being recognised as one of the hallmarks of cancer, cancer-promoting inflammation is a promising target for cancer therapy^1^. Among various processes causing cancer-promoting inflammation, the pathway of cyclooxygenase 2 (COX2) and its metabolite prostaglandin E_2_ (PGE_2_) has been widely accepted as important in human cancers^2^. COXs are rate-limiting enzymes required for the biosynthesis of PGs in the arachidonic acid cascade, and PGE_2_ is the most abundant COX metabolite. Under physiological conditions, COX2 and PGE_2_ are induced during inflammatory processes and act as pro-inflammatory factors^3^. COX2 overexpression and subsequent PGE_2_ overproduction are observed in various human cancers^4–7^ and play a crucial role in the development of the tumour-promoting inflammatory microenvironment^2,8^.

Canine urothelial carcinoma (cUC) is the most common malignancy affecting the lower urinary tract of dogs. cUC is a unique tumour often well managed using COX inhibitors or non-steroidal anti-inflammatory drugs (NSAIDs)^9^. In addition, COX2 is overexpressed in cUC^10–12^. We previously reported that cUC cell lines overexpress PGE_2_ *in vitro* compared to other canine tumour cell lines with different tissues of origin^13^. Further, we suggested that aberrant PGE_2_ production is important for development of the tumour microenvironment and not for cell proliferation or survival^13^. However, the pathway that induces upregulation of COX2/PGE_2_ axis in cUC cells was not elucidated.

Another characteristic of cUC is that a single nucleotide mutation in the *BRAF* gene, V595E, is detected in 70%–80% of canine patients^14,15^. BRAF is an isoform of RAF serine/threonine kinase, which belongs to the RAF/MEK/ERK pathway. This pathway is one of the most important signalling pathways that transmit extracellular signals to cell nuclei, thereby regulating cell proliferation, differentiation, survival, and various other cellular functions. The canine homologous mutation BRAF^V595E^, which is recognized as BRAF^V600E^, is frequently observed in a variety of human malignancies such as malignant melanoma, colorectal cancer, and papillary thyroid cancer^16–18^. The BRAF^V600E^ mutation reportedly induces oncogenic cellular proliferation via constitutive activation of the RAF/MEK/ERK pathway^16,19^. Therefore, several molecular targeting drugs against BRAF^V600E^ have been established and have improved the prognosis of patients with cancer^20,21^. Although canine BRAF^V595E^ is also suggested to contribute to constitutive activation of the RAF/MEK/ERK signalling cascade, its importance in cUC progression remains unclear.

In this study, we screened molecular targeting agents to determine the pathways involved in PGE_2_ production in *BRAF* mutant cUC cells. Subsequently, we investigated the contribution of the RAF/MEK/ERK pathway in the regulation of the COX2/PGE_2_ axis in *BRAF* mutant cUC cells. We also analysed the association between COX2 expression and the *BRAF* genotype in cUC tissues. Our findings indicate a novel association between oncogenic activation of the RAF/MEK/ERK pathway in *BRAF* mutant cUC cells and dysregulation of the COX2/PGE2 axis.

## Results

### In vitro drug screening for disruption of PGE_2_ production in *BRAF* mutant cUC cells

We previously reported that cUC cell lines overexpress PGE_2_^13^. To elucidate the mechanisms underlying aberrant PGE_2_ production in cUC cells, we screened 332 inhibitor compounds using SCADS inhibitor kit 1-4 obtained from Molecular Profiling Committee, Grant-in-Aid for Scientific Research on Innovative Areas “Advanced Animal Model Support (AdAMS)” from The Ministry of Education, Culture, Sports, Science and Technology, Japan (KAKENHI 16H06276; see Supplementary Table S1). A *BRAF* mutant cUC cell line, Sora, was treated with each inhibitor compound at 10 µM for 12 h. The amount of PGE_2_ in the medium was quantified after the treatment, and percent change in PGE_2_ production with respect to that in vehicle control (DMSO) was calculated (Fig. 1A and see Supplementary Table S1). Eighty compounds showed ≥50% reduction in PGE_2_ production in cUC cells. After categorization of all the compounds into their specific target biological pathways, enrichment of each category for the PGE2-suppressing compounds was analysed. Statistical analysis revealed that compounds targeting the arachidonic acid cascade (FDR = 0.086), RAF/MEK/ERK pathway (FDR = 0.067), and p38/JNK pathway (FDR = 0.067) were enriched in these 80 compounds (Table 1 and Fig. 1A and 1B). In addition, the compounds against the enriched pathways did not show strong cytotoxic effects on *BRAF* mutant cUC cells (Fig. 1B). Since the arachidonic acid cascade falls directly upstream of PGE_2_ production, we focused on the RAF/MEK/ERK and p38/JNK pathways as potential biological activators of the COX2/PGE_2_ axis in cUC cells.

**Table 1.**
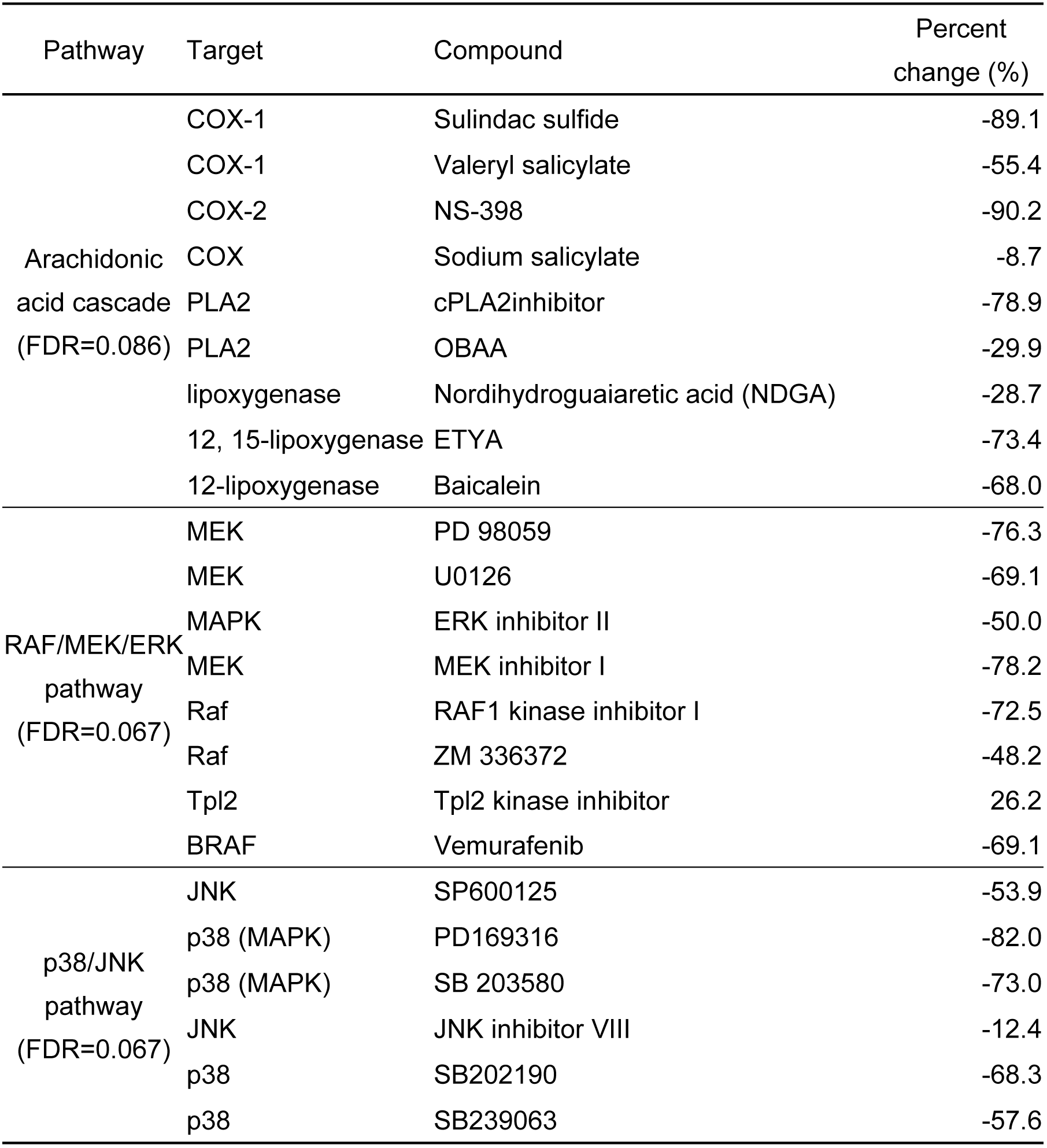
Percent change in PGE2 by the treatment of the inhibitors targeting Arachidonic acid cascade, RAF/MEK/ERK pathway, and p38/JNK pathway.

**Figure 1.**
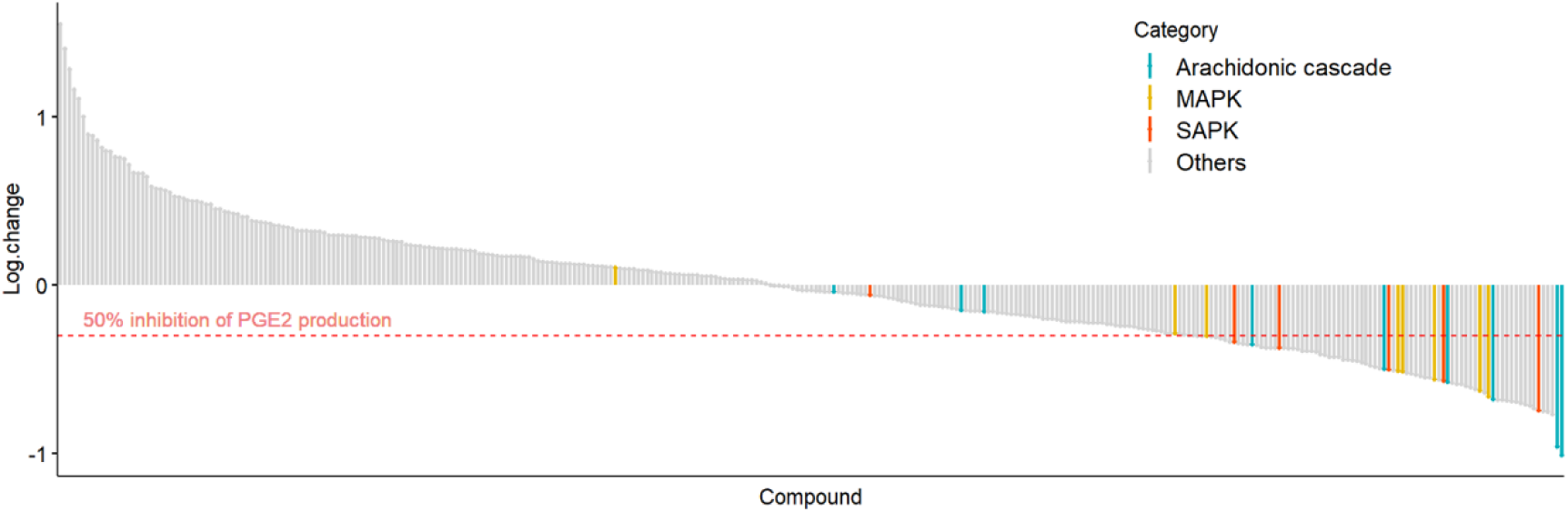
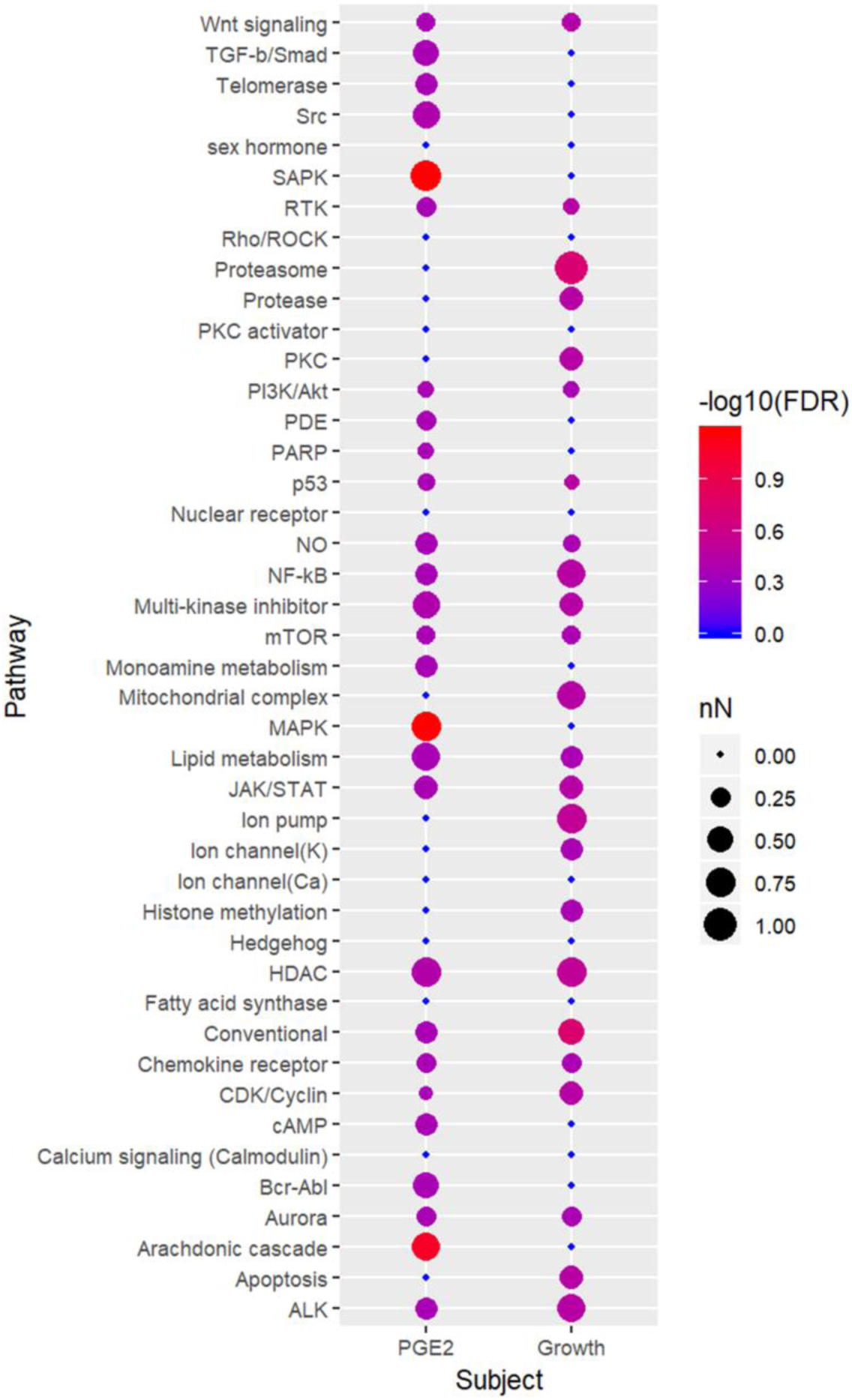
PGE_2_ production in cUC cells (Sora) during drug screening. (A) Y axis represents log 10 values of changes in PGE_2_ production for each inhibitor (n = 332). The drugs which belong to selected pathways are coloured as indicated. (B) Visualization and comparison of the results from drug screening for PGE2 production and cell growth inhibition_42_. Drugs which showed ≥50% reduction in PGE_2_ production and ≥80% reduction in cell viability were included in statistical analysis. Categories which have more than 3 inhibitors are presented in this graph. Node size represents proportion of the drugs which showed reduction more than the threshold in each category. Colour represents statistical significance.

### Effects of MAPK inhibition on expression of the COX2/PGE_2_ axis

To evaluate whether the RAF/MEK/ERK and p38/JNK pathways are involved in PGE_2_ production in *BRAF* mutant cUC cells, Sora cells were treated with the following compounds that specifically inhibit components of various pathways: dabrafenib (BRAF inhibitor), LY3009120 (pan-RAF inhibitor), PD0325901 (MEK inhibitor), SCH772984 (ERK inhibitor), SB239063 (p38 inhibitor), and SP600125 (JNK inhibitor). All compounds were used at 1 µM for 0, 6, 12, and 24 h. After 6 h of treatment, we noted that COX2 expression and PGE_2_ production were decreased by RAF/MEK/ERK inhibitors (Fig. 2A). In the case of the p38/JNK pathway, although p38 inhibition led to the suppression of COX2 expression and PGE_2_ production, JNK inhibition increased the expression of COX2 despite decreased PGE_2_ production (see Supplementary Fig. S1A, B). Further, the cells were exposed to each compound at varying concentrations for 12 h. In this case too, RAF/MEK/ERK inhibitors suppressed COX2 expression and PGE_2_ production in cUC cells in a dose-dependent manner (Fig. 2B and see Supplementary Fig. S1B). Finally, six *BRAF* mutant cUC cell lines (TCCUB, Love, Nene, NMTCC, LTCC, and MCTCC) and one *BRAF* wild type cUC cell line (OMTCC) were treated with the inhibitors with the varying doses for 12 h. In five of the seven cell lines, which showed comparatively higher PGE_2_ production when cultured in the basal medium (see Supplementary Fig. S2), decreases in COX2 expression and PGE_2_ production were observed when the RAF/MEK/ERK pathway was inhibited (Fig. 2C). On the contrary, OMTCC and MCTCC, which show mild expression of the COX2/PGE_2_ axis among the seven cUC cell lines, did not show change in COX2/PGE_2_ expression (Fig. 2C).

**Figure 2.**
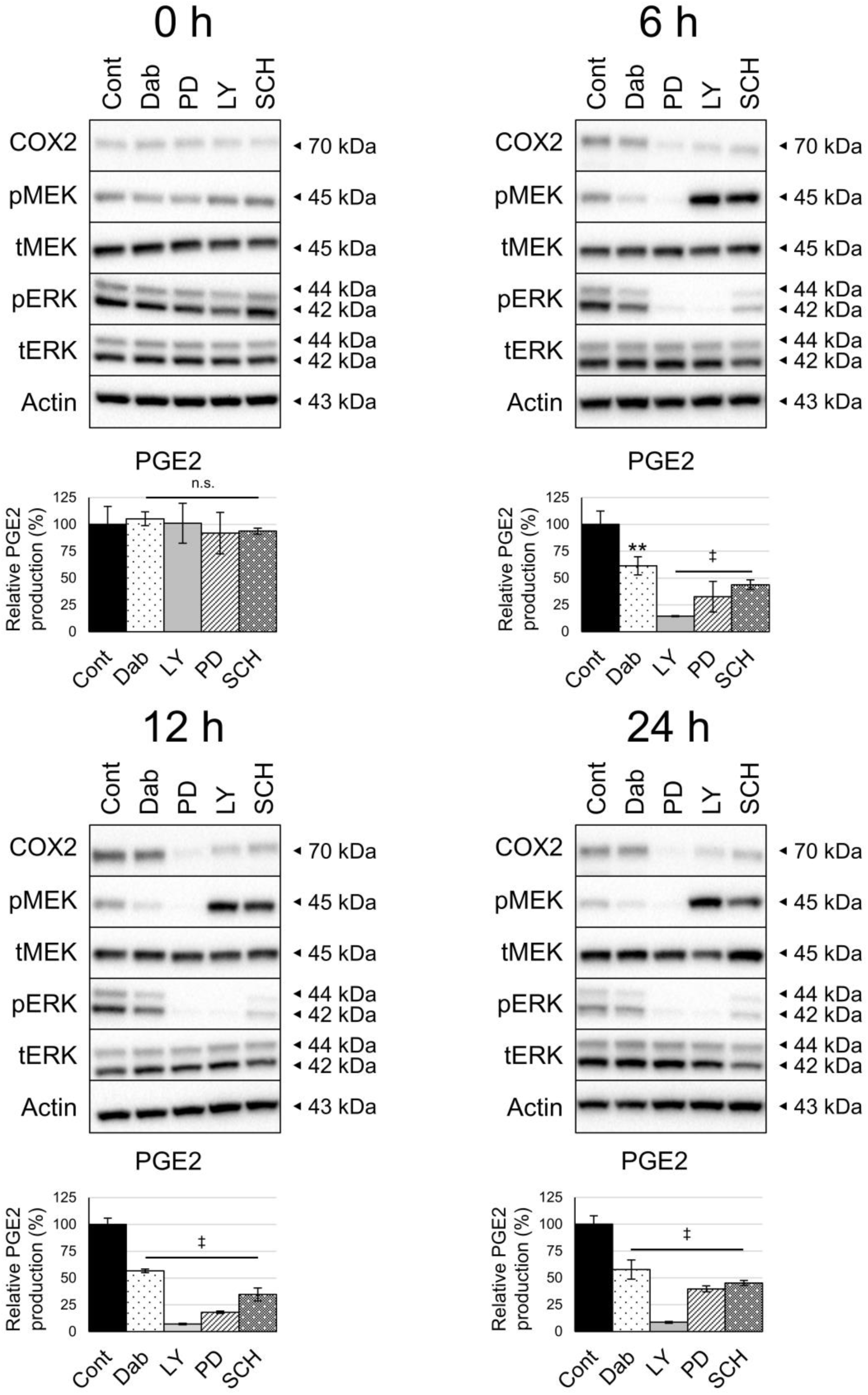
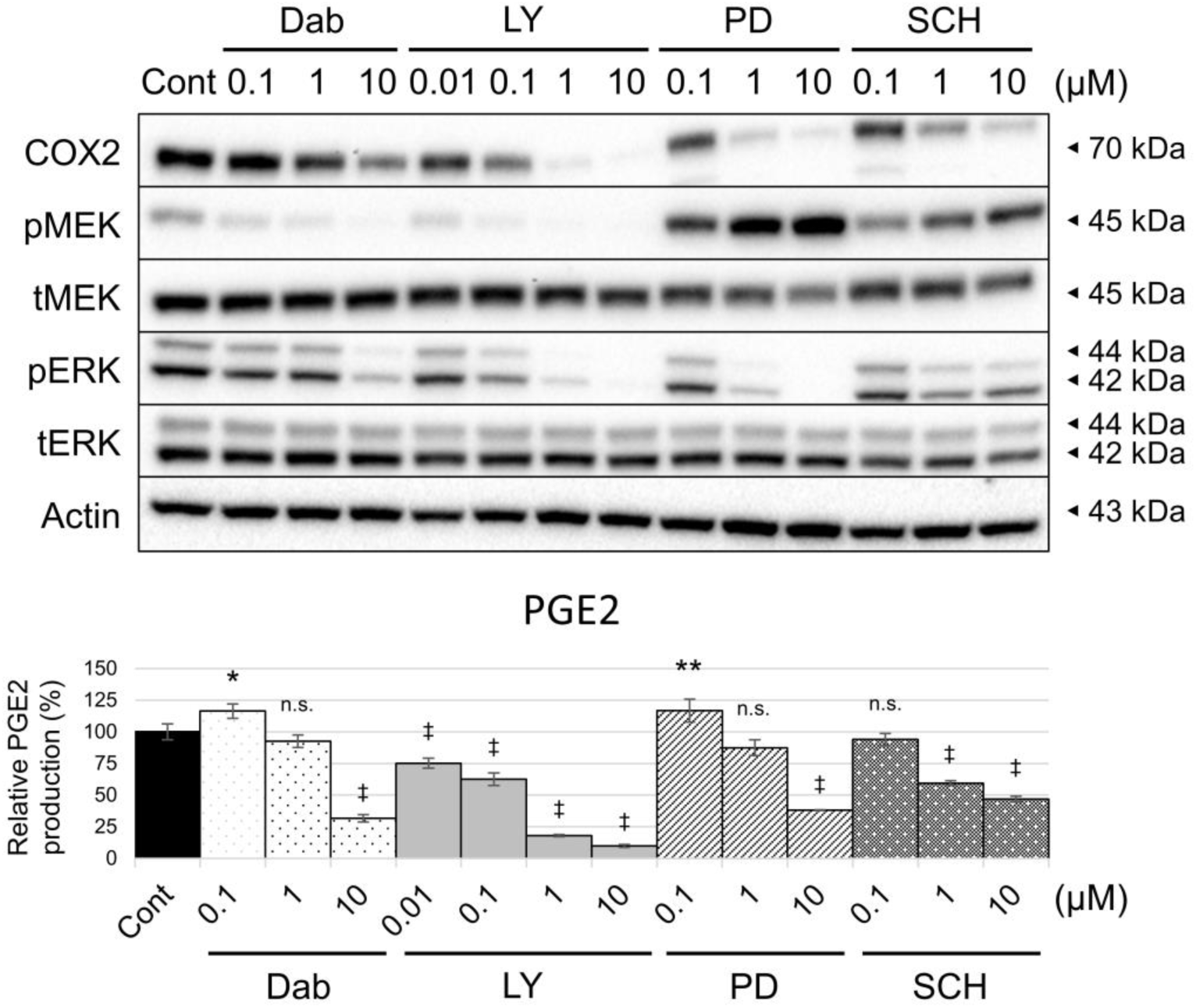
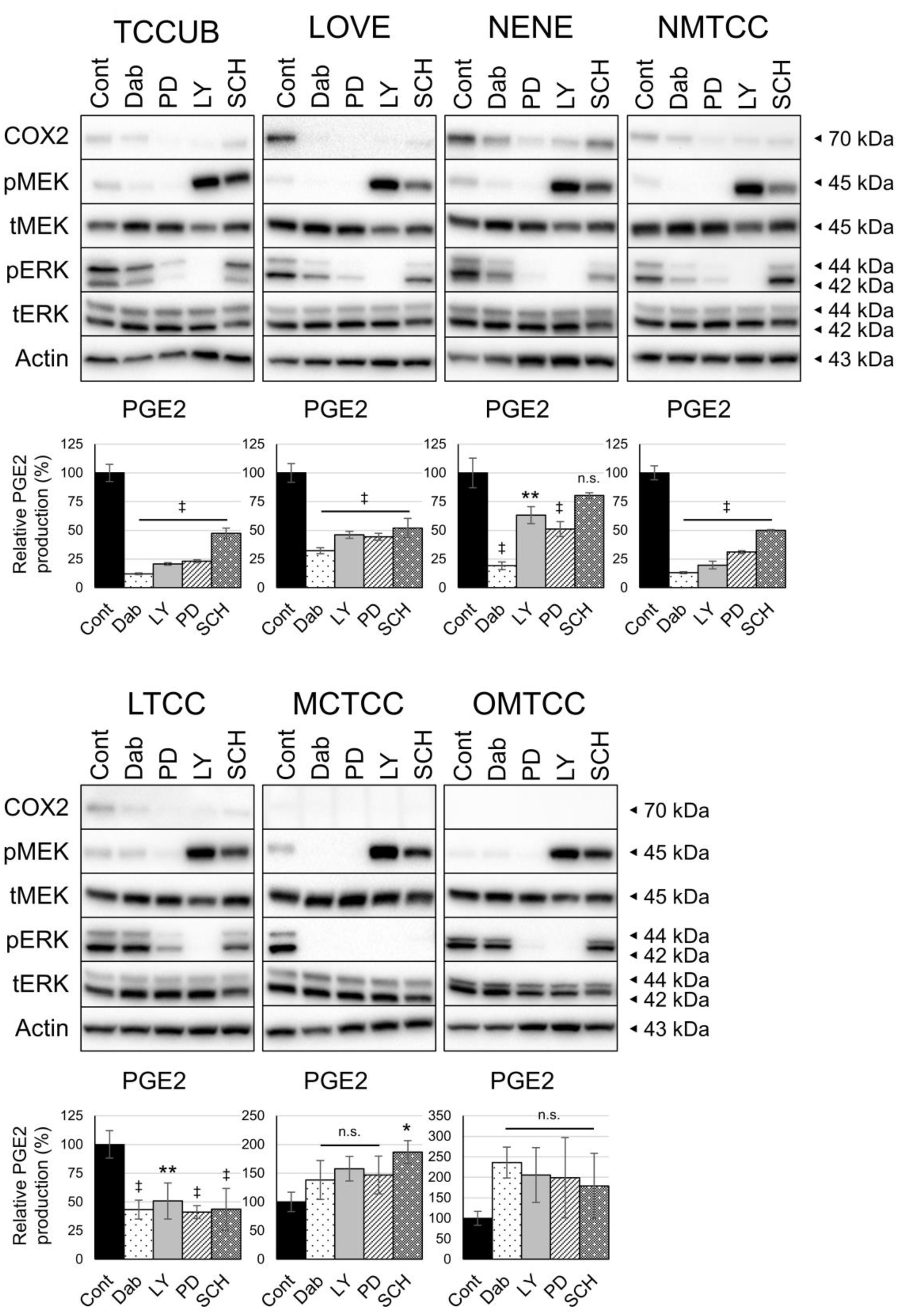
Effect of RAF/MEK/ERK inhibition on COX2 expression and PGE_2_ production. Protein levels in whole cell lysates were detected by Western blotting with Actin as loading control. Amount of PGE_2_ in culture medium was measured by enzyme-linked immunosorbent assay and normalized to cell number. Bar graph represents % control of PGE_2_ production. (A) cUC cells (Sora) were treated with vehicle (dimethyl sulfoxide; Cont) and inhibitors of BRAF (Dabrafenib; Dab), pan-RAF (LY3009120; LY), MEK (PD0325901; PD), and ERK (SCH772984; SCH) at 1 μM for indicated time. (B) cUC cells (Sora) was treated with vehicle (Cont), Dabrafenib (Dab), LY3009120 (LY), PD0325901 (PD), and SCH772984 (SCH) for 12 h at indicated dose. (C) cUC cell lines (TCCUB, Love, Nene, NMTCC, LTCC, MCTCC, and OMTCC) were treated with vehicle (Cont), Dabrafenib (Dab) at 10 μM, LY3009120 (LY) at 1 μM, PD0325901 (PD) at 1 μM, and SCH772984 (SCH) at 1 μM for 12 h. Data are presented as mean ± SD of three experiments. * indicates p < 0.05, ** p < 0.01, ‡ p < 0.001 compared to vehicle control (Dunnett’s test).

### Association between *BRAF* mutation and COX2 expression

To further examine the association between the COX2/PGE_2_ axis and the RAF/MEK/ERK pathway in patients with cUC, we performed immunohistochemical analysis of COX2 expression and genotyping of the *BRAF* gene. The levels of COX2 expression were analysed using a semi-quantitative method described in previous reports (Fig. 3a)^10,22,23^. Digital PCR-based *BRAF* genotyping identified *BRAF* mutation in 33 cUC tissues, while 10 tumours were *BRAF* wild-type. cUC tissues with *BRAF* mutation tended to express higher levels of COX2, although the difference between mutant tumours and wild-type tumours was not significant (p = 0.0569; Fig. 3b and 3c)

**Figure 3.**
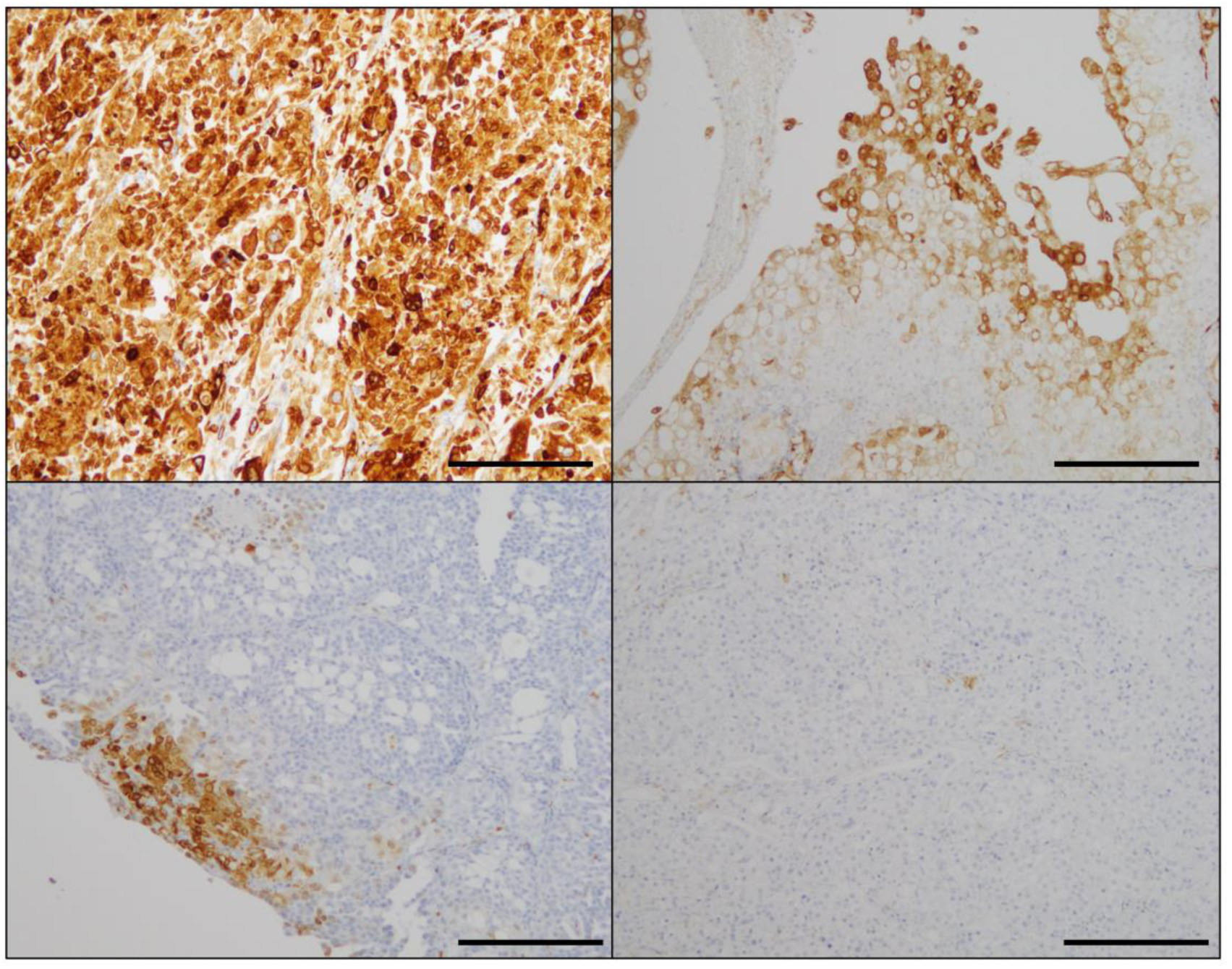
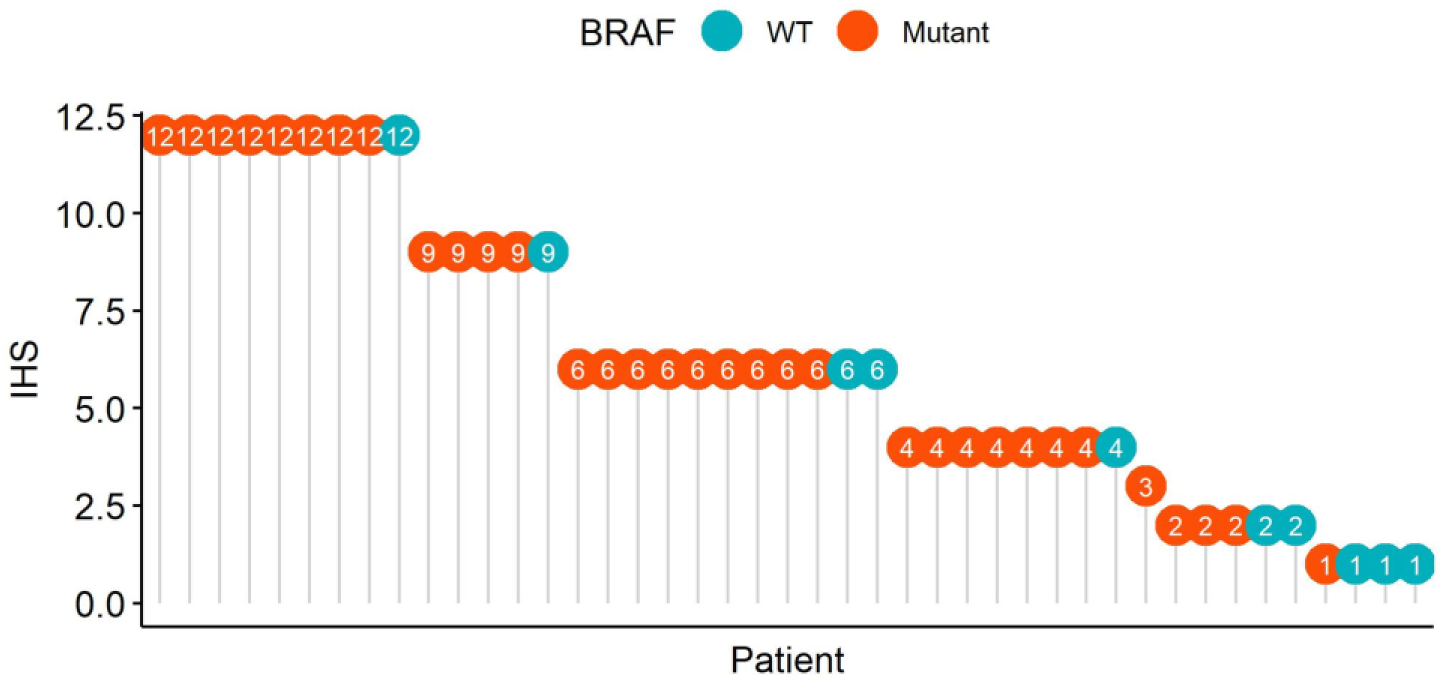
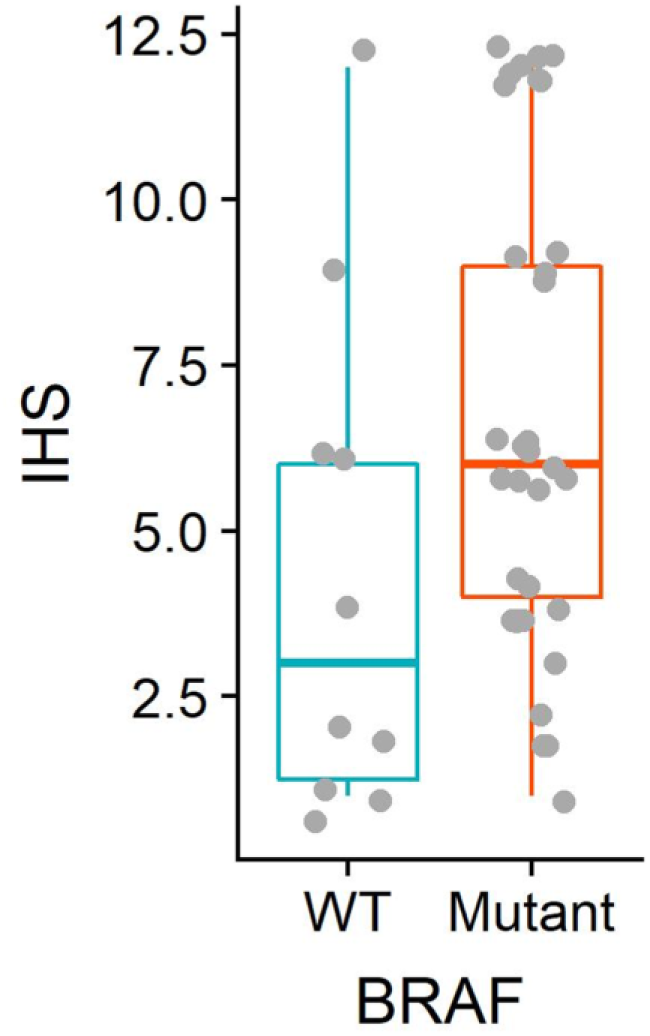
Levels of COX2 expression in cUC tissue. (A) Representative image for each immunohistochemical score (IHS) at 100-fold magnification. IHS was determined by a semi-quantitative method as described in Materials and Methods. IHSs of presented pictures were 12 (top left), 9 (top right), 6 (bottom left), 2 (bottom right), respectively. Scale bar: 200 μm. (B) A lollipop chart displays COX2 scores for individual patients and their relationship with BRAF genotype. (C) A summarizing box plot for COX2 expression in BRAF wild-type (WT; red) and mutant cUC tissue (Mutant; blue). Y axis represents IHS. p = 0.0569 (Wilcoxon exact test).

## Discussion

The COX2/PGE_2_ axis plays a crucial role in tumour development. In several human tumours, it has been revealed that the COX2/PGE_2_ axis is affected by oncogenic mutations^24,25^. A few recent reports suggested that BRAF^V600E^, one of the most common driver mutations in human cancer, contributed to COX2/PGE_2_ overexpression^26^. Our results of drug screening indicated that among various biological signalling pathways, the RAF/MEK/ERK pathway facilitates PGE_2_ production in cUC cells. We further showed that oncogenic activation of the RAF/MEK/ERK pathway may lead to aberrant upregulation of the COX2/PGE_2_ axis in *BRAF* mutant cUC cells. We also suggested that the *BRAF* genotype in cUC tissues may predict overexpression of COX2. From these observations, we concluded that highly penetrated *BRAF* mutation in cUC cells probably contributes to activation of the RAF/MEK/ERK pathway and subsequent induction of the COX2/PGE_2_ axis to prepare the pro-tumoral microenvironment. This, in turn, may confer notable COX inhibitor vulnerability to the tumour, as widely observed in veterinary clinical practice.

Recent studies have reported that activation of the RAF/MEK/ERK pathway promotes COX2 expression and PGE_2_ production in human cancers. A couple of studies suggested that ERK signalling activated by hepatocyte growth factor leads to increased COX2 expression in human colorectal cancer and non-small cell lung cancer ^27,28^. Further, a recent study that investigated *BRAF* mutant melanoma cells established from genetically engineered mice suggested that COX2/PGE_2_ expression depends on RAF/MEK/ERK pathway activation^26^. In this study, we suggested that the activation of the RAF/MEK/ERK pathway is involved in the overexpression of the COX2/PGE_2_ axis in *BRAF* mutant cUC cells. Thus, our findings indicate close analogy of biological mechanisms of carcinogenesis or tumour progression between humans and dogs.

cUC is a unique tumour often managed with NSAID monotherapy, indicating the importance of the COX2/PGE_2_ axis in the development and progression of cUC^9^. Since no direct cytotoxicity was observed in our previous study^13^, the therapeutic effect of NSAIDs was considered to be exerted via blocking the effect of eicosanoids, specifically PGE_2_, on tumour microenvironment. This hypothesis was further supported in this study by the fact that no correlation was observed between suppression of PGE2 production and cell growth. Therefore, the findings of this study suggest that *BRAF* mutation might render cUC cells capable of developing their pro-tumoral microenvironment. Although this principle should be further investigated in *in vivo* model, cUC would serve as a valuable model to reveal the association between oncogenic activation of the RAF/MEK/ERK pathway and development of the pro-tumoral microenvironment via the COX2/PGE_2_ axis.

In this study, the BRAF inhibitor dabrafenib showed weaker inhibitory effect on ERK phosphorylation and COX2/PGE_2_ expression than other inhibitors. In addition, during drug screening, all inhibitors targeting the RAF/MEK/ERK pathway did not show strong inhibition of cUC cell growth even at high concentration. Several kinds of human cancers also show similar primary resistance with ERK reactivation to BRAF inhibitors^29–31^. These instances of primary resistance are attributed to “paradoxical activation” of ERK signalling through other RAF isoforms or RAF dimers^30,32,33^. The resistance of cUC tot BRAF inhibitors observed in this study may have the same underlying mechanisms, since MEK or pan-RAF inhibitors were able to suppress ERK phosphorylation for a long period.

PGE_2_, one of the products of COX2, has been reported to be an inducer of cancer-promoting inflammation^2^. On the contrary, in the arachidonic acid cascade, there are some tumour suppressors such as PGD2^34,35^. In our previous study, *BRAF* mutant cUC cells also produced large amounts of PGD2 compared to other cells^13^. Although in the current study, we focused on PGE_2_ as it is a major product of COX2, PGD2 or other tumour-suppressing factors could be regulated by the activation of the RAF/MEK/ERK pathway in BRAF mutant cUC cells. The regulation of other products and the overall effect of RAF/MEK/ERK inhibition on tumour microenvironment should be further investigated.

In vitro drug screening revealed the candidate mechanisms for increased production of PGE_2_ in cUC cells. Although we focused on the RAF/MEK/ERK pathway in this study, the stress-activated protein kinases p38 and JNK were also suggested to be important for PGE_2_ overproduction in initial screening. p38 and JNK are members of the MAPK family and activated by environmental stress, including ultraviolet exposure, oxidative stress, or inflammatory cytokines^36^. They have been suggested to induce COX2 expression in inflammatory processes, such as rheumatoid arthritis or Parkinson’s disease, and in human cancer-promoting inflammation^37–39^. Our results suggested that the p38/JNK pathway also regulates PGE_2_ production in cUC cells in a manner different from that of the RAF/MEK/ERK pathway. Therefore, the mechanisms underlying the activation of the p38/JNK pathway and its role in cUC progression requires further investigation.

In the histological analysis, *BRAF* mutant cUC cells tended to show higher expression of COX2. In a recent study, the association between *BRAF* mutation and COX2 expression was only observed in cUC tissues of specific dog breeds ^40^. Further, in our *in vitro* study, one of the *BRAF* mutant cUC cell lines did not overexpress COX2 and PGE_2_. Notably, this cell line did not show ERK reactivation upon BRAF inhibition, indicating that *BRAF* mutation does not necessarily cause RAF/MEK/ERK pathway activation in the same manner. Another possibility is that the RAF/MEK/ERK pathway is activated by different mutations in *BRAF* wildtype tumours. Further study about dysregulation of the RAF/MEK/ERK pathway as well as variability in the consequences of *BRAF* mutations will lead to further understanding of cUC pathogenesis.

In conclusion, our study revealed that overexpression of the COX2/PGE_2_ axis in cUC cells harbouring the *BRAF* mutation is related to the activation of the RAF/MEK/ERK pathway. The importance of COX2/PGE2 expression and BRAF mutation has been highlighted in several human cancers. Therefore, our findings can be extrapolated to human medicine as well. In the future, validation through “gain of function” studies using genetic engineering techniques accompanied by *in vivo* studies is required. Moreover, a greater patient sample size for immunohistochemical analysis would be beneficial.

cUC is a unique tumour that shows notable response to COX inhibitors. We believe that cUC can be an excellent model to study the relationship between oncogenic mutations and establishment of pro-tumoral microenvironment. Further, our data suggested the existence of other potent regulatory mechanisms for COX2/PGE_2_ expression. Further investigation of this characteristic of cancer will provide new insights into the association of driver mutations and cancer-promoting inflammation in humans and animals.

## Methods

### Cell culture

Seven cUC cell lines, TCCUB, Sora, Love, Nene (originally established in our laboratory), NMTCC, MCTCC, and OMTCC (kindly provided by Hokkaido University) were used^13,41,42^. Each cell line was maintained in RPMI-1640 supplemented with 10% heat-inactivated foetal bovine serum (FBS) and 5 mg/L gentamicin at 37° C in a humidified atmosphere with 5% CO_2_.

### Drugs

For in vitro drug screening, the SCADS inhibitor kits (kit 1, ver 3.3; kit 2, ver 2.0; kit 3, ver 1.6; kit 4, ver 2.3) were kindly provided by Molecular Profiling Committee, Grant-in-Aid for Scientific Research on Innovative Areas “Platform of Advanced Animal Model Support” from The Ministry of Education, Culture, Sports, Science and Technology, Japan (KAKENHI 16H06276). For the following experiment, dabrafenib (BRAF inhibitor), PD0325901 (MEK inhibitor), and SCH772984 (ERK inhibitor) were purchased from Selleck. LY3009120 (pan-RAF inhibitor) was purchased from Cayman Chemical. These inhibitors were reconstituted in DMSO and stored at −20° C or −80° C.

### PGE_2_ measurement

PGE_2_ concentration in culture supernatant was measured using enzyme-linked immunosorbent assay kit (Cayman Chemical) according to the manufacturer’s instructions. To normalize the amount of PGE_2_ to the cell density, direct cell count or sulforhodamine B (SRB) assay was performed^43^.

### In vitro drug screening

Sora cells were seeded in a 96-well plate at a density of 11,000 cells per well. After 24-h incubation, the culture medium in each well was discarded, and fresh medium containing the drugs from the SCADS inhibitor kits was added at a final concentration of 10 µM. After 12-h incubation, the culture supernatants were collected. After normalizing PGE_2_ production to cell density as mentioned above, percentage change in PGE_2_ production from control was calculated.

### Growth inhibition

To compare with the changes in PGE2 production, data regarding cell growth inhibition after treatment with the inhibitors from the SCADS inhibitor kits were obtained from our previous study. Detailed protocol is described in our previous study^24^. Briefly, Sora cells were treated with the drugs from the SCADS inhibitor kits at 10 µM as described in *In vitro drug screening* section. After 72-h incubation, cell densities were determined by SRB assay.

### Protein detection

cUC cells were seeded in serum-free medium and incubated for 24 h as in our previous report (see Supplementary Table S2)^24^. After 24-h serum starvation, cUC cells were treated with final concentration of 10% FBS and RAF/MEK/ERK inhibitors as indicated in each figure. After treatment for the indicated times, cells were lysed for 30 min on ice in RIPA buffer consisting of 50 mM Tris-HCl, 150 mM NaCl, 5 mM EDTA, 0.1% sodium dodecyl sulphate, 1% Triton-X, 10 mM NaF, 2 mM NaVO_4_, and complete protease inhibitor cocktail (Roche Diagnostics). After centrifugation, protein concentrations were measured using bicinchoninic acid protein assay kit. Equal amounts (10 µg) of total protein were separated by sodium dodecyl sulphate-polyacrylamide gel electrophoresis and transferred to polyvinylidene difluoride membranes. The membranes were blocked with 5 % skim milk in Tris-buffered saline containing 0.1% Tween (TBST) for 1 h at room temperature to avoid non-specific antibody binding. Further, the membranes were incubated overnight at 4° C with each primary antibody (see Supplementary Table S3). The membranes were then washed with TBST and incubated with horseradish peroxidase-conjugated anti-mouse or anti-rabbit secondary antibody (1:10000) from GE Healthcare for 1 h at room temperature. Protein signal was developed with a chemiluminescent system (Merck Millipore) and captured with an imaging system (BioRad Laboratories).

### Immunohistochemistry

Formalin-fixed paraffin-embedded (FFPE) cUC tissues were retrospectively evaluated. Surgically resected cUC tissue samples (n = 43) were obtained from the archival collection of the Veterinary Medical Center of the University of Tokyo (samples collected from 2009 to 2012 and 2015 to 2016) and Veterinary Medical Teaching Hospital of Nippon Veterinary and Life Science University (from 2003 to 2017). The clients from respective hospitals provided informed consent for the use of these samples for this study. cUC tissues of 4 μm thickness were deparaffinized in xylene and hydrated in graded alcohol. Antigen retrieval was performed for 10 min at 121° C in citrate buffer (pH 6.0). The sections were then treated with 3% hydrogen peroxide for 30 min at room temperature. After blocking with 5% normal goat serum in TBST for 1 h at room temperature, the sections were incubated overnight with 1:100 of anti-COX2 mouse monoclonal antibody (BD Bioscience) at 4° C in a humidified chamber. Subsequently, the sections were rinsed with TBST and then incubated with EnVision polymer reagent for mouse (Dako) for 60 min at room temperature. After rinsing with TBST again, peroxidase reactions were developed for 3 min with 3,3′-diaminobenzidine (Dako). The sections were counterstained with haematoxylin.

### BRAF genotyping

*BRAF* genotype of cUC tissue was determined as described in a previous study^42^. Briefly, gDNA was extracted from FFPE sections of cUC tissue using Qiaamp DNA FFPE Tissue Kit (Qiagen). BRAF^V595E^ and wild-type *BRAF* gene were amplified with Taqman probe and signal detection was performed using QuantStudio 3D Digital PCR System (Thermo Fisher Scientific).

### Immunohistochemical evaluation

For scoring COX2 expression in cUC tissue, a semi-quantitative immunohistochemical score (IHS) system described in a previous report was used^10,22,23^. Briefly, the percentage of COX2-positive tumour cells in the entire section and their signal intensity were evaluated when viewed at 100x and 200x magnification, respectively. Positivity was graded as 1 = <1%, 2 = 1%–9%, 3 = 10%–50%, 4 = >50%, and intensity was graded as 0 = no staining, 1 = mild staining, 2 = moderate staining, 3 = strong staining. IHS was obtained by multiplying positivity score and intensity score.

### Statistical analysis

Statistical analysis was performed using R software (https://www.R-project.org/). To determine enriched pathways revealed in drug screening, all of the 331 compounds were divided into 98 groups according to their targeting pathways, and p value for each pathway was calculated using hypergeometric distribution. Multiple comparison was adjusted by Benjamini–Hochberg method and the FDR cut-off was set as 0.1. To test the effect of RAF/MEK/ERK inhibition on PGE_2_ production, Dunnett’s multiple comparison test was used. Difference in the IHS score depending on the *BRAF* genotype was calculated using Wilcoxon rank sum test. All values are shown as the mean value ± standard error of the mean. P values < 0.05 were considered statistically significant.

## Acknowledgements

We thank Molecular Profiling Committee, Grant-in-Aid for Scientific Research on Innovative Areas “Platform of Advanced Animal Model Support” from the Ministry of Education, Culture, Sports, Science and Technology, Japan, which is supported by KAKENHI 16H06276 for providing the drug screening kits. Also, we are grateful to Yuki Hoshino (Iwate University) for providing the cell lines (NMTCC, LTCC, MCTCC, and OMTCC) and Masaki Michishita (Nippon Veterinary and Life Science University) for providing the FFPE cUC tissues. This work was supported by JSPS KAKENHI Grant Number 16K20968 and 18K14582 (KS).

## Author Contributions

RY, KS, RN, and TN designed this study. RY performed the experiments. SE and MS contributed to the data analysis. RY and KS prepared the figures and wrote this manuscript. KS, RN, HS, YE, NF, RN, and TN supervised this study.

## Data Availability

The data obtained during the current study will be available from the corresponding author upon reasonable request.

