## Supplementary Figures for "Aberrant expression of the COX2/PGE_2_ axis is induced by activation of the RAF/MEK/ERK pathway in BRAF^V595E^ canine urothelial carcinoma"

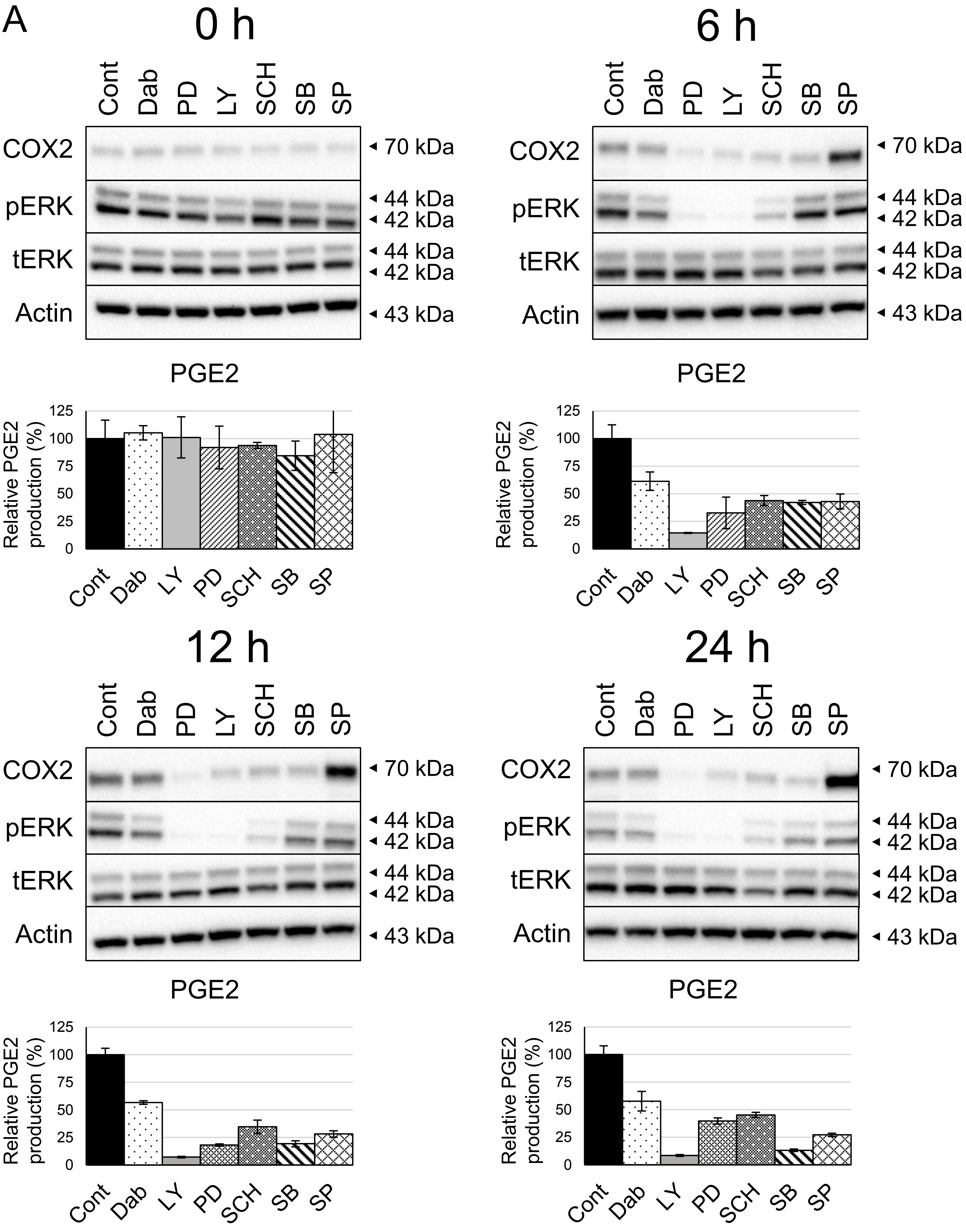
A
0 h
Cont
Dab
PD
LY
SCH
SB
SP
COX2
pERK
tERK
Actin
70 kDa
44 kDa
42 kDa
44 kDa
42 kDa
43 kDa
PGE2
Relative PGE2 production (%)
125
100
75
50
25
0
Cont
Dab
LY
PD
SCH
SB
SP
6 h
12 h
24 h

B

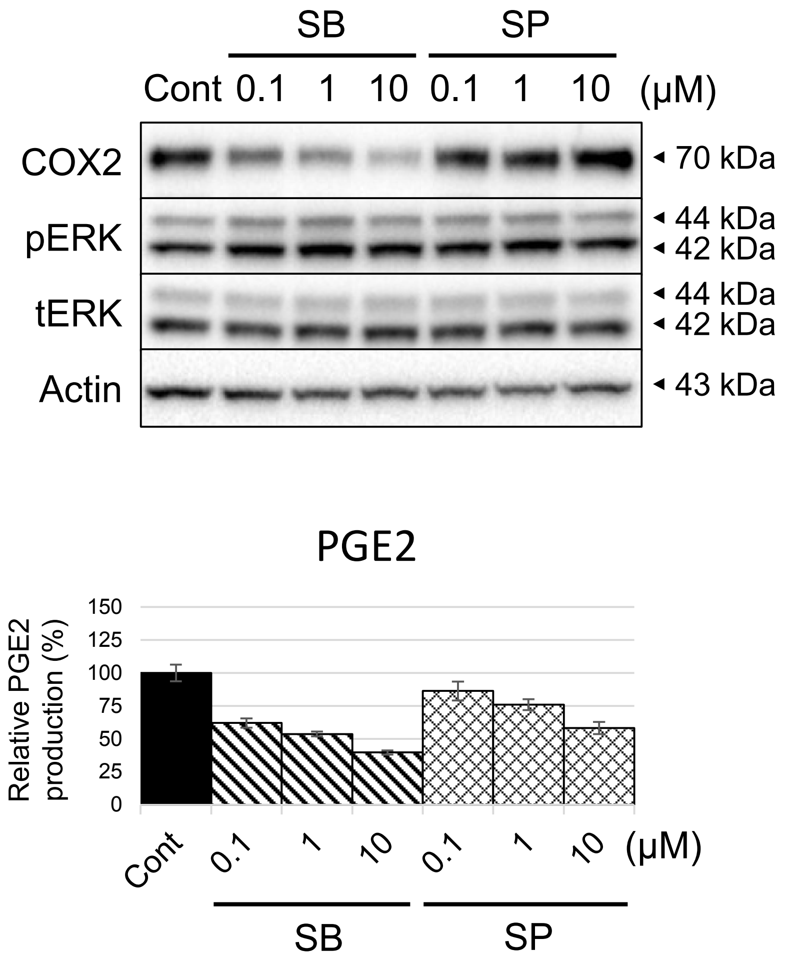


**Supplementary Figure S1**. Effect of p38/JNK pathway inhibition on COX2 expression and PGE2 production. Protein levels in whole cell lysate were detected by Western Blotting with Actin as loading control. Amount of PGE2 in culture medium were measured by enzyme linked immunosorbent assay and corrected to cell number. Bar graph represents % control of PGE2 production. (A) Whole image of the membrane and PGE2 production shown in Figure. 1A. UC cells (Sora) was treated with vehicle (dimethyl sulfoxide; Cont) and inhibitors of BRAF (Dabrafenib; Dab), pan-RAF (LY3009120; LY), MEK (PD0325901; PD), ERK (SCH772984; SCH), p38 (SB239063; SB), and JNK (SP600125; SP) at 1 μM for indicated time. (B) cUC cells (Sora) was treated with vehicle (Cont), SB239063 (SB), and SP600125 (SB) for 12 h at indicated dose. Data are presented as mean ± SD of three experiments.

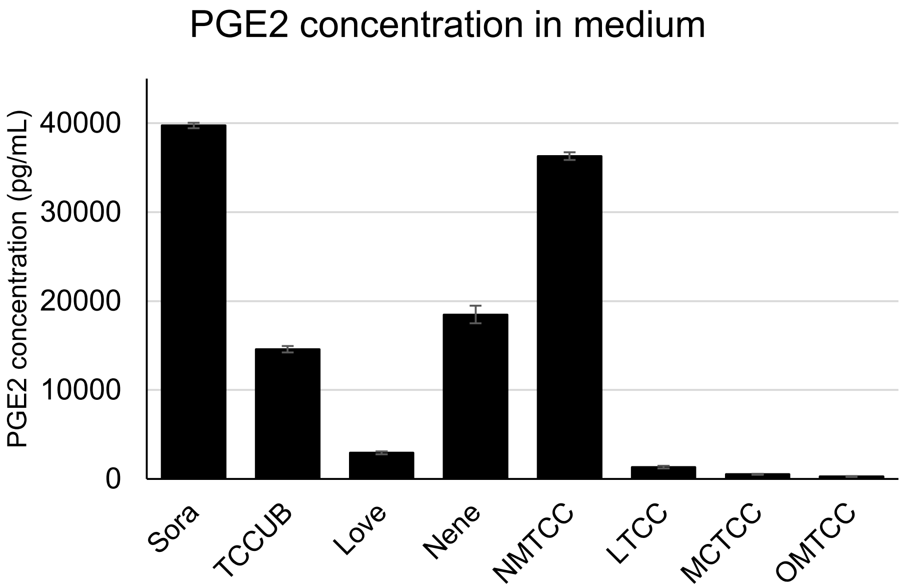


**Supplementary Figure S2**. Basal PGE2 production of cUC cell lines. cUC cell lines (Sora, TCCUB, Love, Nene, LTCC, MCTCC, and OMTCC) were seeded in serum free medium and incubated for 24 h. After 24 h serum starvation, cUC cells were treated with final concentration of 10% Fatal bovine serum and incubated for further 24 h. PGE2 concentration in culture supernatant was measured using enzume-linked immunosorbent assay. Data are presented as mean ± SD of three experiments.
